# Blocking Formation of Neurotoxic Reactive Astrocytes is Beneficial Following Stroke

**DOI:** 10.1101/2023.10.11.561918

**Authors:** Kimberly Prescott, Alexandra E. Münch, Evan Brahms, Maya M. Weigel, Kenya Inoue, Marion S. Buckwalter, Shane A. Liddelow, Todd C. Peterson

## Abstract

Microglia and astrocytes play an important role in the neuroinflammatory response and contribute to both the destruction of neighboring tissue as well as the resolution of inflammation following stroke. These reactive glial cells are highly heterogeneous at both the transcriptomic and functional level. Depending upon the stimulus, microglia and astrocytes mount a complex, and specific response composed of distinct microglial and astrocyte substates. These substates ultimately drive the landscape of the initiation and recovery from the adverse stimulus. In one state, inflammation- and damage-induced microglia release tumor necrosis factor (TNF), interleukin 1α (IL1α), and complement component 1q (C1q), together ‘TIC’. This cocktail of cytokines drives astrocytes into a neurotoxic reactive astrocyte (nRA) substate. This nRA substate is associated with loss of many physiological astrocyte functions (e.g., synapse formation and maturation, phagocytosis, among others), as well as a gain-of-function release of neurotoxic long-chain fatty acids which kill neighboring cells. Here we report that transgenic removal of TIC led to reduction of gliosis, infarct expansion, and worsened functional deficits in the acute and delayed stages following stroke. Our results suggest that TIC cytokines, and likely nRAs play an important role that may maintain neuroinflammation and inhibit functional motor recovery after ischemic stroke. This is the first report that this paradigm is relevant in stroke and that therapies against nRAs may be a novel means to treat patients. Since nRAs are evolutionarily conserved from rodents to humans and present in multiple neurodegenerative diseases and injuries, further identification of mechanistic role of nRAs will lead to a better understanding of the neuroinflammatory response and the development of new therapies.

## Blocking Formation of Neurotoxic Reactive Astrocytes is Beneficial Following Stroke

Stroke is the second-leading cause of death worldwide, and one in four people who reach the age of 25 will have a stroke in their lifetime (Feigin et al., 2022). An ischemic stroke occurs when blood supply to a region of the brain ceases due to blockage of a blood vessel (usually by formation of a blood clot) and is one of the most common types of strokes. Current stroke therapies include intravenous and endovascular thrombolysis, and mechanical clot removal, but treatment only helps patients whose stroke has not completed prior to arriving at the hospital and endovascular therapies are not available at most hospitals. Current FDA approved treatments for ischemic stroke aim at returning blood flow to the affected area by breaking up the blood clot, with the only approved drug being recombinant tissue plasminogen activator (rt-PA). While the use of rt-PA is beneficial in stroke outcomes for some patients, there is a very small therapeutic window, with only 2% or less of ischemic stroke patients eligible to receive rt-PA (Tseng et al., 2020; Wardlaw et al., 2012; Xiong et al., 2022).

Initial cell death occurs within minutes of the onset of stroke, and these dead and dying cells make up the ischemic infarct core. Additional brain cells in the surrounding tissue, known as the peri-infarct area, are vulnerable during the acute and sub-acute period and their death leads to infarct expansion (Lakhan et al., 2009). Many pathophysiological mechanisms including oxidative stress, excitotoxity, hypoxia due to brain edema, and neuroinflammation all contribute to this cell death in the subacute period (Khoshnam et al., 2017; Lakhan et al., 2009; Pantcheva et al., 2014).

The neuroinflammatory response is complex and has both beneficial and detrimental attributes related to stroke recovery. In the CNS, the inflammatory response is primarily orchestrated by the release of cytokines from glial cells, such as microglia and astrocytes (Jayaraj et al., 2019; Sun et al., 2015). During post-stroke neuroinflammation, microglia alter their morphology, increase phagocytosis, and alter the types and levels cytokines they secrete (Stence et al., 2001; Zhang, 2019). During acute stages of ischemia, microglia may release anti-inflammatory cytokines, with a maintenance of proinflammatory cytokine release as the inflammatory response progresses (Hu et al., 2012). We and others have previously demonstrated that tumor necrosis factor (TNF), interleukin 1α (IL1α), and complement component 1q (C1q), together ‘TIC’, are released by microglia in the peri-infarct region following stroke, and are required and sufficient to transform astrocytes into a neurotoxic substate (Hu et al., 2012; Liddelow et al., 2017).

These neurotoxic reactive astrocytes (nRA) lose many normal physiological functions including their ability to induce and support synapses, reuptake and recycle glutamate, and downregulate phagocytosis pathways (Barbar et al., 2020; Guttenplan et al., 2021; Liddelow & Barres, 2017; Liddelow et al., 2017). On the other hand, nRAs have an intriguing gain-of-function ability, namely the release of fully saturated long chain free fatty acids which are sufficient to induce caspase-3 mediated lipoapoptotic neuron death (Guttenplan et al., 2021) . In the context of stroke, this lipid-mediated cell death is potentially responsible for further expanding the infarct area.

To examine the role that these nRAs play in ischemic stroke, transgenic, triple knockout (TKO) mice unable to produce the TIC cytokines can be used. We previously used these TKO mice to determine that the release of the TIC cytokines by microglia are important in transforming astrocytes into the neurotoxic substate. In a mouse model of glaucoma, TKO mice uncovered that TIC cytokine-induced nRAs contributed to retinal ganglion cell neuron death (Guttenplan et al., 2020a; Sterling et al., 2020). The inhibition of microglia-derived TIC cytokines has also been achieved pharmacologically using a glucagon-like 1 peptide receptor agonist in the pre-formed alpha synuclein mouse model of Parkinson’s disease. Here block of microglia-induced nRA reactivity was sufficient to decrease neuronal cell loss and improve motor outcomes (Yun et al., 2018). Given these promising results, and the localization of nRAs following stroke, we tested here whether TKO mice would have reduced astrogliosis, decreased infarct size, and improved motor outcomes after ischemic stroke compared to wild type (WT) mice.

## Materials and Methods

### Animals

Seventy-six female C57BL/6J mice (WT; Jackson Laboratory, Ellsworth, ME) and female transgenic mice (*TNF^-/-^, IL1-α^-/-^ and C1q^-/-^*; TKO) at approximately 8 weeks of age were used. Mice had *ad libitum* access to food and water, and were housed up to 5 animals per cage. Mice were housed at room temperature (22 ± 1 °C) on a 12-hour reverse light/dark cycle. All experimental procedures and protocols were approved by the Stanford University Institutional Animal Care and Use committee (IACUC).

### Study Design

To characterize the pathophysiological differences after stroke, the distal middle cerebral artery occlusion (dMCAO) model was used in the acute and subacute time points. Mice underwent permanent focal ischemia using the dMCAO model. WT (N = 15) and TKO (N = 16) mice were perfused 24 hours after dMCAO to examine the acute phase, and WT (N = 10) and TKO (N = 8) mice were perfused 7 days after dMCAO to examine the subacute phase (**Fig. 1a**). While the dMCAO model is a more clinically relevant model for middle cerebral artery occlusion known for highly reproduceable cortical lesions, it does not typically produce long-term behavioral deficits in mice (Doyle & Buckwalter, 2014; Doyle et al., 2012; Peterson et al., 2021). To characterize the differences during the chronic time point and test motor outcomes, WT (N = 16) and TKO (N = 11) mice underwent a photothrombotic stroke over the motor cortex and behavior was assessed. Prior to sacrifice, behavior assessments were performed prior to and 1-, 3-, 7-, 14-, and 28 days post-stroke. We assessed gross motor movement using the rotating beam task and fine motor movement was assessed using the tapered beam task and foot fault task. Animals were perfused with 0.1 M phosphate buffered saline 30 days after stroke and prepared for tissue processing and immunohistochemistry.

**Figure 1.**
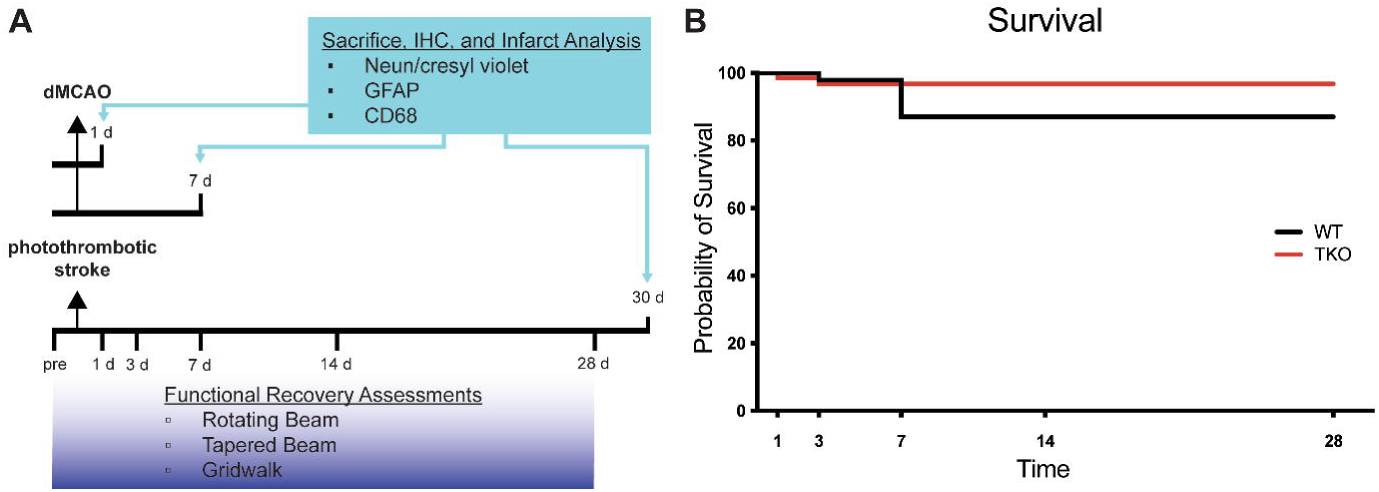
Experimental timeline and survival rate of animals 30 days after photothrombotic stroke. **(A)** Timeline showing 1- and 7-day sacrifice after dMCAO and 30-day sacrifice following photothrombotic stroke. Immunohistochemistry was performed to examine infarct volume (NeuN/cresyl violet), astrogliosis (GFAP) and microgliosis (CD68). Functional recovery assessment was performed up to 28 days after photothrombotic stroke include the rotating beam, tapered beam, and grid walk tasks. **(B)** Survival rates were not significantly different between wildtype (WT) and *Tnf^-/-^Il1a^-/-^C1qa^-/-^* triple knockout (TKO) mice after photothrombotic stroke (*X^2^* = 1.85, *p* = .17). Abbreviations: IHC; immunohistochemistry, GFAP; glial fibrillary acidic protein, CD68; Cluster of Differentiation 68, d, day; dMCAO, distal middle cerebral artery occlusion; TKO, *Tnf^-/-^Il1a^-/-^C1qa^-/-^* triple knockout mouse; WT, wild type.

### Stroke Models

Mice were anesthetized using 2% Isoflurane in 2 L/min oxygen and temperature was maintained at 37 °C. For the 24 hour and 7-day sacrifice timepoints, mice underwent a dMCAO to create a primary cortical infarct as described (Doyle & Buckwalter, 2014; Peterson et al., 2021). Briefly, an incision was made in the skin over the temporalis muscle to expose the medial cerebral artery through the skull. A craniotomy was performed, and the medial cerebral artery was then cauterized. The temporalis muscle and skin were replaced, and the incision closed. Buprenorphine (0.1 mg/kg) and Cefazolin (25 mg/kg) were injected subcutaneously.

To assess chronic inflammation and functional recovery, the photothrombotic stroke model was performed targeting the motor cortex. Mice were injected intraperitonially with Rose Bengal (10 mg/ml in sterile saline; 40 mg/kg of body weight), anesthetized, and then mounted on a stereotactic frame. A Metal Halide Fiber optic light (no. 56371, Edmund Optics) with a 1 mm diameter was directly placed over the right motor cortex (-0.5 mm anterior, +1.5 mm lateral to bregma) for 15 min. The incision was closed and a subcutaneous injection of Buprenorphine (0.1 mg/kg) and Cefazolin (25 mg/kg) was administered.

### Functional Recovery Assessment

Motor behavior tasks were performed in WT and TKO mice 1-, 3-, 7-, 14-, and 28 days after photothrombotic surgery and included the rotating beam, tapered beam, and foot fault tasks described previously (Cheng et al., 2016; Cheng et al., 2014; Peterson et al., 2021; Schaar et al., 2010). For all tasks, there was one habituation day and two pre-training days that occurred prior to stroke induction. For the rotating beam task, animals crossed a rotating beam (6 rpm) longitudinally and the distance covered before falling was recorded for four trials per session. For the tapered beam test, the number of contralateral foot faults made as they progressed down the beam for four trials per session was recorded. For the foot fault task, animals were allowed to freely roam for 2 min and the number of contralateral forelimb faults was recorded.

### Tissue Collection

Mice were anesthetized with 87.5 mg/kg Ketamine/12.5 mg/kg Xylazine and transcardially perfused with 10 U/mL heparin in 0.9% NaCl before the brain was dissected and drop fixed in 4% paraformaldehyde in phosphate buffer overnight. The brain was then transferred to 30% sucrose in phosphate buffered saline (PBS). Coronal sections (40 μm) were sectioned using a freezing sliding microtome (Microm HM430), and sequentially placed in 16 tubes filled with cryoprotective medium (30% glycerin, 30% ethylene glycol, 40% 0.5 M sodium phosphate buffer) and stored at 20 °C until processing.

### Immunohistochemistry

Free floating tissue was stained using standard immunohistochemistry procedures. Briefly, sections were washed and agitated in 3% serum (donkey, EMD Millipore, Cat# 566460, rabbit, Vector, Cat# S-5000; goat, Vector, Cat# S-1000) prior to antibody incubation. Primary antibodies (NeuN, 1:20,000, Millipore, MAB377B; CD68, 1:1,000, Serotec, MCA1957S; GFAP, 1:10,000, Dako Z0334) were diluted in 0.3% Triton X in PBS and incubated for 12 hours at RT. After washing in PBS, secondary antibodies (NeuN n/a biotinylated primary; anti-rabbit, 1:500, Vector, BA-1000; anti-rat, 1:500, Boster, BA1058) were incubated for 60 minutes at RT. For cresyl violet counter staining, mounted tissue was rehydrated before being stained in cresyl violet for 5 min and dehydrated through graduated ethanol before cover slipping with entellan mounting medium.

### Quantification Procedures

To measure stroke volume, 10 coronal sections spaced exactly 320 μm apart were stained with NeuN, and then counter stained with cresyl violet (see above). Sections were imaged using a PathScan Enabler IV (Meyer Instruments, Housten, TX) and the infarct and hemisphere volumes traced using Fiji (version 2.9.0/1.53t, (Schindelin et al., 2012). To control for edema on the side ipsilateral of stroke that commonly occurs in the dMCAO model, the percent of infarcted hemisphere (infarcted area (contralateral hemisphere area / (ipsilateral area without infarct + infarct area)) / contralateral hemisphere X 100) was used (Nouraee et al., 2019; Peterson et al., 2021). The photothrombotic stroke model has little edema associated with it 29 days after stroke, so the direct infarct volume was used for measurements.

To image macrophages and astrocytes, five sections spaced 640 μm apart were immunostained with CD68 and GFAP; respectively. Two images were taken (Keyence microscope) at 40x magnification in the peri-infarct cortex, one medial and one lateral to the stroke core. Each image was one view field (543 μm) from the stroke border. Fiji was used to convert the images to 8-bit color (black and white) and blindly quantified images at a uniform threshold encompassing the area covered (density) of the soma and processes using Fiji (Peterson et al., 2021).

### Statistical Analyses

GraphPad Prism (Software, 2023) and RStudio (Team, 2020) were used to perform graphing and statistical analysis. Significance was determined with a *p* < .05 and values are expressed as the mean ± standard error. Survival data was analyzed using the Kaplan-Meier method. Immunohistochemical data were analyzed using a one-tailed Student’s *t* test for 2 groups. For the behavior tasks, a linear mixed effects model was analyzed using RStudio and the lme4 R package (Bates et al., 2015). Mice were specified as a random factor to help account for unequal group sizes due to mortality. A one-way analysis of variance (ANOVA) was performed to test the differences between models using the lmerTest package (Kuznetsova et al., 2017). Where differences were found, a Dunnett’s post-hoc test was performed to examine differences by day.

## Results

### Global Deletion of *Tnf*, *IL1*α, and *C1q* Does Not Alter Survival Following Stroke

Survival rates of animals following ischemia can be indicative of stroke severity. To test if TKO animals had altered survival compared to WT animals, we used the photothrombotic model of stroke and tracked 26 animals out to 30 days. We found no difference in the survival of WT and TKO mice over this time period, (*p* = .174, **Fig. 1b**). There were also no significant differences in mortality in the acute and subacute time period following dMCAO (data not shown). Our data suggests that while TIC cytokines play important roles in inflammation, the removal of these pro-inflammatory cytokines does not increase the chance of surviving following stroke.

### Global Deletion of *Tnf, IL1*α, and *C1q* Decreased Astrocyte Cytoskeletal Restructuring After Stroke

To examine how TIC cytokines influence the neuroinflammatory response after stroke, the area covered by both macrophages and astrocytes in TKO and WT mice was measured. To assess astrocyte cytoskeletal structure, we immunohistostained for GFAP in the peri-infarct area during the acute (24 hours), subacute (7 days), and chronic (30 days) phases after stroke. There was no difference in the amount of GFAP immunoreactivity 24 hours post-stroke between the TKO and WT mice (*p* = .104, **Fig. 2b**), however TKO mice had significantly reduced GFAP levels compared to the WT mice 7 days post-stroke (*p* = .011) and 30 days post-stroke (*p* < .0001; **Fig. 2c, d**). While many have reported that alterations in GFAP levels alone are indicative of changes in astrocyte reactivity, we would caution this interpretation from this data alone.

**Figure 2.**
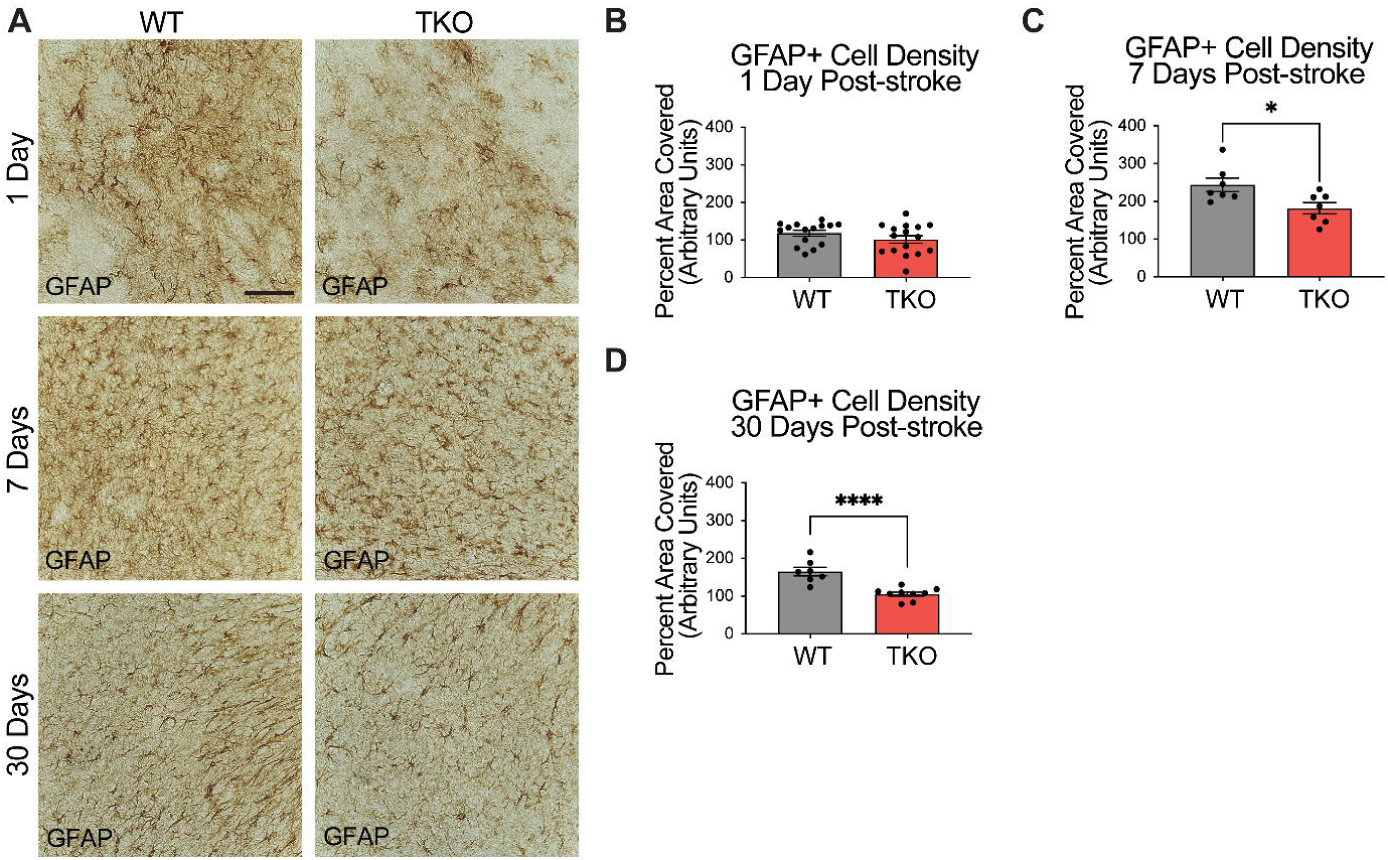
TNF, IL1α, and C1q knockout reduces GFAP+ cell number in the peri-infarct area 7- and 30-days post ischemic stroke. **(A)** Representative images of GFAP+ cells in the peri-infarct area for wildtype (WT) and *Tnf^-/-^Il1a^-/-^C1qa^-/-^*triple knockout (TKO) mice 1-, 7-, and 30-days post ischemia. **(B)** No difference was found in GFAP+ cells in the peri-infarct area 1 day after ischemia between the WT and TKO mice, t (29) = 1.29, *p* = .104. **(C)** TKO mice had significantly reduced GFAP+ cells in the peri-infarct area than WT mice 7 days post-stroke, t (12) = 2.64, *p* = .011. **(D)** TKO mice had significantly reduced GFAP+ cells in the peri-infarct area than WT mice 30 days post-stroke, t (14) = 5.03, *p* < .0001. * *p* < .05, ** *p* < .001, *** *p* < .0001. Abbreviations: GFAP; glial fibrillary acidic protein, TKO, *Tnf^-/-^Il1a^-/-^C1qa^-/-^* triple knockout mouse; WT, wild type.

### *TNF*, *IL1*α, and *C1q* Are Required for Myeloid Cell Upregulation of CD68 Following Stroke

To examine how TIC cytokines collectively influence myeloid cells following ischemic stroke, we used CD68+ immunostaining and quantified cell numbers in the peri-infarct at 24 hours, 7 days, and 30 days post-stroke. There was no difference in macrophage response, measured by the density of CD68 immunostaining at 24 hours post-stroke between the TKO and WT mice (*p* = .285, **Fig. 3a**), but TKO mice had significantly reduced CD68 immunostaining compared to WT mice 7 days post-stroke (*p* = .001) and 30 days post-stroke (*p* = .0001, **Fig. 3b, c**). Similar to the use of a single marker, GFAP, for astrocyte reactivity, we would caution the interpretation of changes in myeloid cell reactivity states using CD68 alone. Thus, removal of TIC cytokines in the more chronic stages following stroke appears to alter the proinflammatory environment. Not only that, but while the initial upregulation at 24 hours is normal, the TKO mice do not propagate the inflammatory signal 7- and 30 days post-stroke.

**Figure 3.**
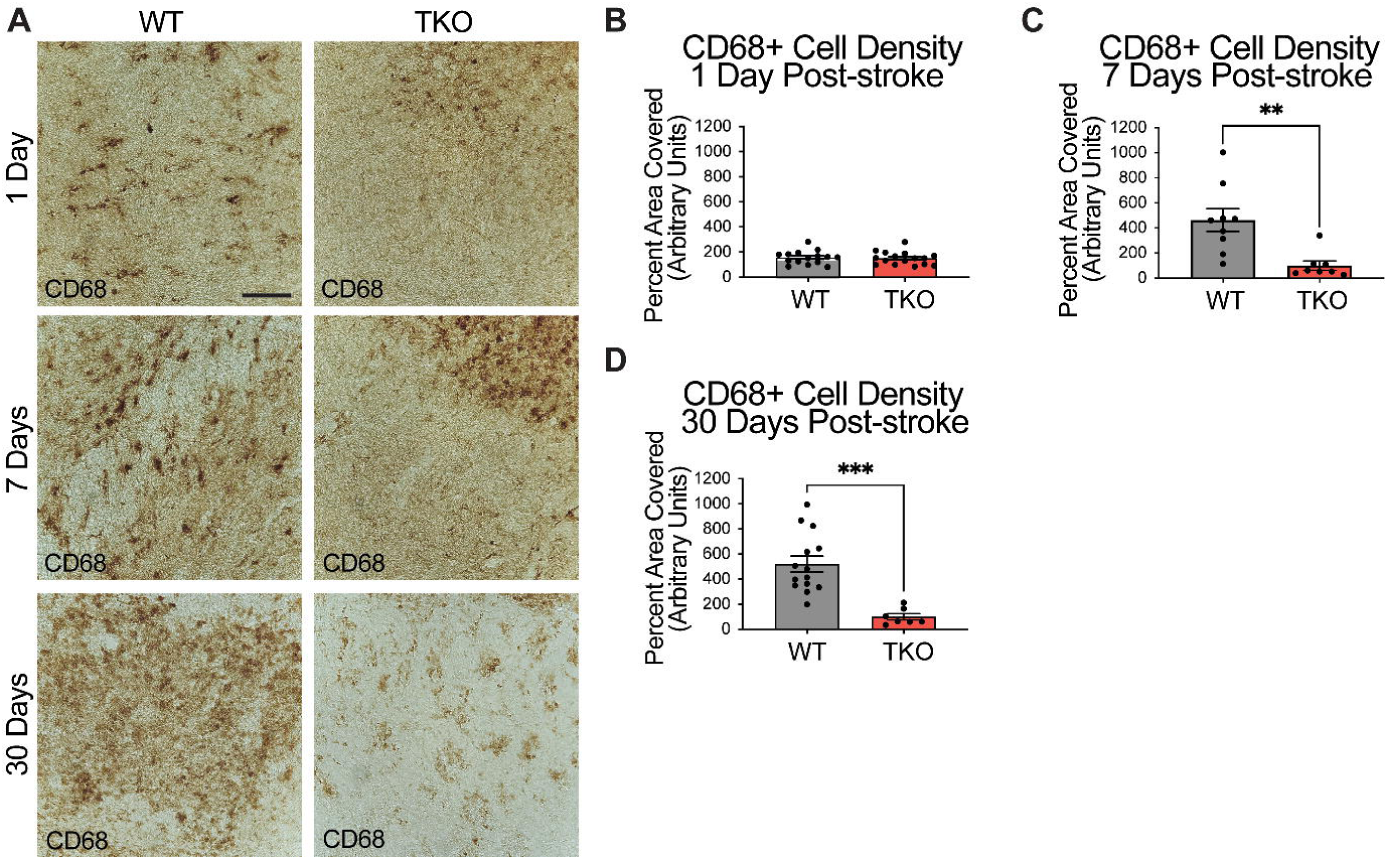
TNF, IL1α, and C1q knockout reduces CD68+ cells in the peri-infarct area 7- and 30-days post ischemic stroke. **(A)** Representative images of the CD68+ cells in the peri-infarct area for wildtype (WT) and *Tnf^-/-^Il1a^-/-^C1qa^-/-^*triple knockout (TKO) mice 1-, 7-, and 30-days post ischemia. **(B)** No difference was found in CD68+ cells in the peri-infarct area 1 day after ischemia between the WT and TKO mice, t (29) = 0.29, *p* = .285. **(C)** TKO mice had significantly reduced CD68+ cells in the peri-infarct area than WT mice 7 days post-stroke, t (15) = 3.51, *p* = .002. **(D)** TKO mice had significantly reduced CD68+ cells in the peri-infarct area than WT mice 30 days post-stroke, t (19) = 4.55, *p* = .0001. * *p* < .05, ** *p* < .001, *** *p* < .0001. Abbreviations: CD68; Cluster of Differentiation 68, TKO, *Tnf^-/-^Il1a^-/-^C1qa^-/-^* triple knockout mouse; WT, wild type.

Together with GFAP staining, these data suggest that in the absence of TIC cytokines, minimal differences occur in astrocyte GFAP and myeloid cell CD68 protein levels immediately following stroke. However, there was a decrease in these astrocyte and myeloid cell markers from 1 week onwards suggesting that the inflammatory response resolves much more quickly in the TKO mice.

### Infarct Volume Following Stroke is Decreased in the Absence of *TNF*, *IL1*α, and *C1q*

Given our discovery of altered astrocyte and myeloid cell responses in the absence of TIC cytokines, we next decided to measure neuron viability in these animals. To assess how TIC cytokines can influence infarct volume during the acute, subacute, and chronic phases after stroke, we used immunohistochemistry of NeuN+ neurons with a cresyl violet counterstain to measure the gross infarct volume. TKO mice had significantly reduced infarct volumes 24 hours (*p* = .002) and 7 days post-stroke (*p* < .0001; **Fig. 4a, b**) using the dMCAO model. During the chronic phase of stroke, TKO mice were also found to have a significantly reduced infarct volume 30 days post photothrombotic stroke of the motor cortex (*p* = .008, **Fig. 4c**). These data suggest that TIC cytokines directly, or TIC cytokine-induced nRAs are consequential to neuron viability following ischemic insult in the brain.

**Figure 4.**
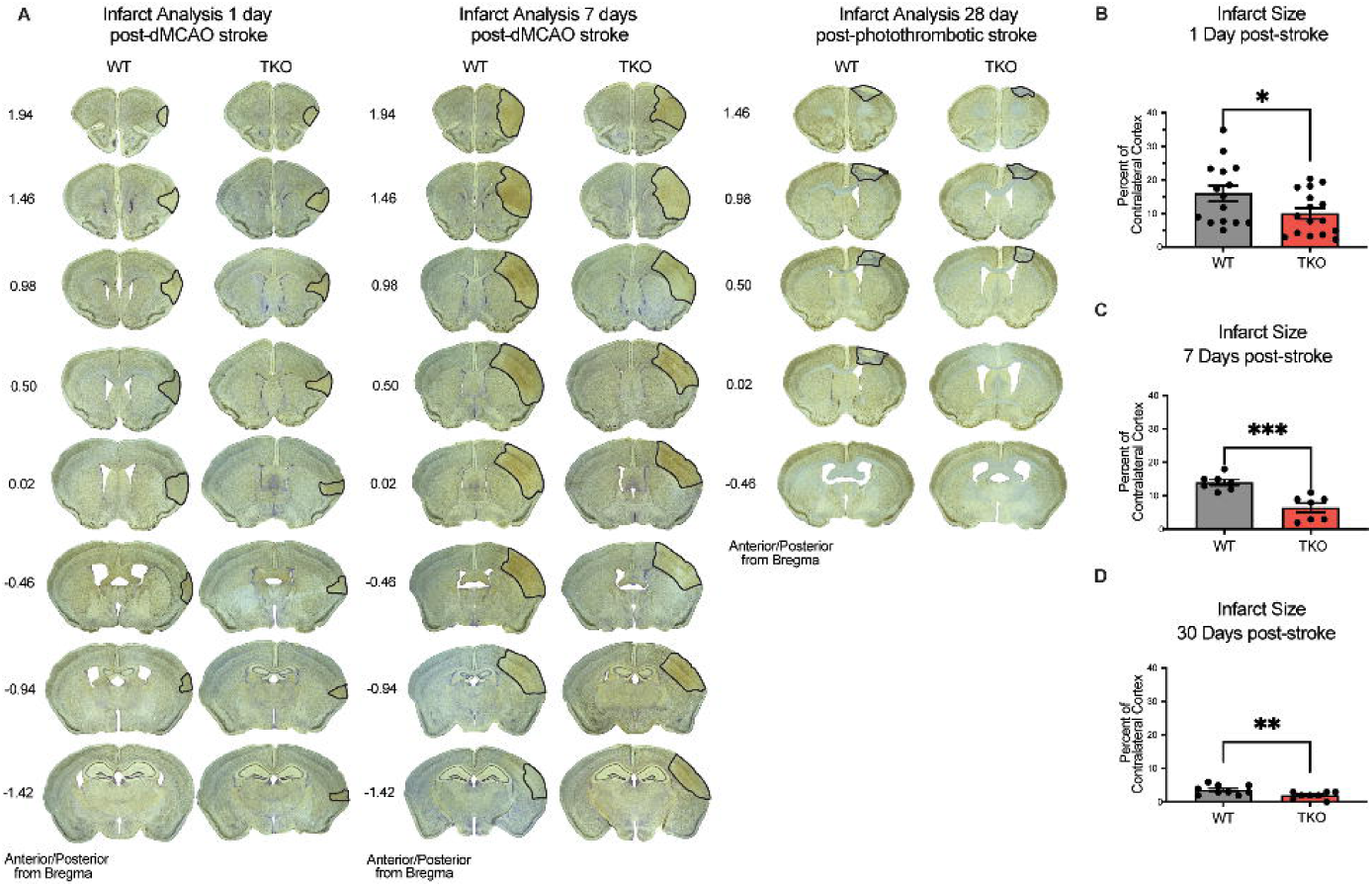
TNF, IL1α, and C1q knockout reduces infract volume 1-, 7-, and 30-days post stroke. **(A)** Representative images of stroke volume using NeuN/cresyl violet stain of wildtype (WT) and *Tnf^-/-^Il1a^-/-^C1qa^-/-^* triple knockout (TKO) mice 1-, 7-, and 30-days post-ischemia. **(B)** TKO mice had significantly reduced infarct volume compared to WT mice 1-day post-stroke, t (29) = 2.09, *p* = .022. **(C)** TKO mice had significantly reduced infarct volume than WT mice 7 days post-stroke, t (12) = 4.64, *p* < .0001. (D) TKO mice had a significantly reduced infarct volume compared to the WT mice 30 days post-stroke, t (15) = 2.712, *p* = .008. * *p* < .05, ** *p* < .001, *** *p* < .0001. Abbreviations: dMCAO, distal middle cerebral artery occlusion; TKO, *Tnf^-/-^ Il1a^-/-^C1qa^-/-^*triple knockout mouse; WT, wild type.

These data suggest that in line with our hypothesis, TIC induced nRAs play a key role in cell death and infarct expansion following stroke. Given these results, we next wanted to determine if this decrease in infarct expansion would correlate with improvements in motor function.

### Functional Deficits After Stroke are Improved in the Absence of *TNF*, *IL1*α, and *C1q*

Since deletion of TIC cytokines improved neuronal viability and decreased infarct volume following dMCAO, we hypothesized that this would lead to improved functional recovery following stroke in the photothrombotic stroke model. We examined motor function using the, a photothrombotic stroke of the motor cortex in adult mice. We quantified limb function contralateral to the stroke 1-, 3-, 7-, 14-, and 28 days post-stroke using a battery of behavioral tests. TKO mice traveled significantly longer on the rotating beam compared to WT mice 1- and 3 days post-stroke (*p* = .026, **Fig.** 5a). TKO mice had significantly fewer contralateral limb errors on the tapered beam compared to WT mice 1-, 3-, 7-, and 14 days post-stroke (*p* = .0189, **Fig. 5b**). This measures fine motor ability. There were no differences of contralateral forelimb foot faults made on the foot fault task between groups (*p* = .91, **Fig. 5a, c**). These data suggest that in addition to decreasing astrocyte cytoskeletal rearrangement, myeloid cell responses, and overall improving neuron viability, deletion of TIC in mice improves their motor function after stroke.

**Figure 5.**
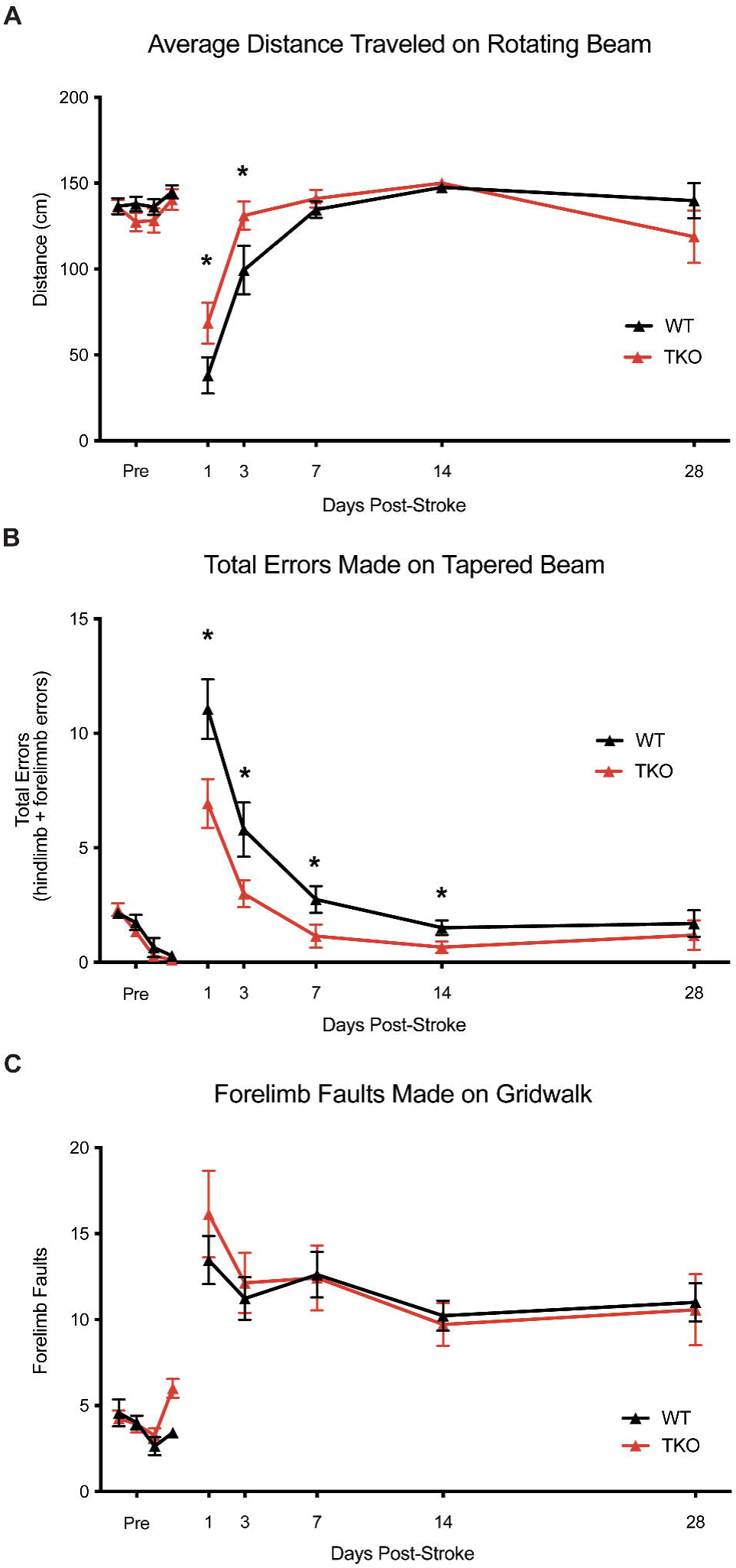
TNF, IL1α, and C1q deletion reduces behavioral deficits after ischemic stroke. **(A)** *Tnf^-/-^Il1a^-/-^C1qa^-/-^* triple knockout (TKO) mice a significantly further distance on the rotating beam as the wildtype (WT) mice, *F* (1, 24) = 0.3969, *p* = .026. A post-hoc analysis revealed that TKO mice made fewer errors 1-(*p* = .0358) and 3 days (*p* = .0244) after stroke. **(B)** TKO mice mad significantly fewer errors on the tapered beam task than WT mice, *F* (1, 24) = 6.342, *p* = .0189. A post-hoc analysis revealed that TKO mice made fewer errors 1-(*p* < .0124), 3- (*p* < .023), 7- (*p* < .0329), and 14 (*p* < .0453) days after stroke. **(C)** TKO mice and WT mice made a similar number of contralateral forelimb errors on the grid walk task, *F* (1, 32.008) = 0.015, *p* = .91. * *p* < .05. Abbreviations: TKO, *Tnf^-/-^Il1a^-/-^C1qa^-/-^* triple knockout mouse; WT, wild type.

## Discussion

We examined the role TIC cytokines in the neuroinflammatory response after stroke by using TKO mice with global deletion of these cytokines. These TKO mice had a reduced astrocyte and myeloid cell response after stroke, which translated to a reduction in infarct volume. TKO mice also had reduced motor behavior deficits when compared to WT mice. Previous research demonstrates that these cytokines play an important role in nRA formation, and our research suggests that nRAs also likely play a detrimental role after ischemic stroke (Guttenplan et al., 2020b; Liddelow et al., 2017; Yun et al., 2018). Inhibiting nRAs by targeting the release of these specific cytokines could lead to new stroke therapies aimed at reducing secondary damage caused by the neuroinflammatory response.

We were able to demonstrate a robust reduction in stroke size, which may be due to a direct effect on neurons resulting in less inflammation, or a reduction of neuroinflammation, which caused less neuronal injury. While we were not able to uncouple direct effects of TIC cytokines directly acting on neurons versus nRA functions, the reduction in infarct volume of TKO mice from 24 hours to 30 days remains both exciting and intriguing. TIC cytokines are primarily being produced by reactive microglia (Hernandez et al., 2023; Liddelow et al., 2017) following stroke, and could have direct actions of their own, on neurons for example, in addition to creating nRAs. These effects could be net positive or negative. TNF is largely thought of as a negative factor, but can also have beneficial effects in stroke (Pan & Kastin, 2007). TNF, IL-1, and C1q also play a dual role in regulating synaptic plasticity (Erion, 2014; Liu et al., 2017; Nemeth & Quan, 2021; Steinmetz & Turrigiano, 2010; Stellwagen & Malenka, 2006; Stevens et al., 2007) and may be involved with synaptic reorganization following acute stroke and trauma (Liguz-Lecznar & Kossut, 2013; O’Connor, 2013; Park & Bowers, 2010; Rizzo et al., 2018). However, since TIC cytokines are necessary and sufficient to induce nRAs (Liddelow et al., 2017), it is most likely that TKO mice are reducing the number and magnitude of nRAs, which is leading to this protective effect (Guttenplan et al., 2021; Yun et al., 2018).

Importantly, not all astrocytes after stroke will be neurotoxic, and they are unlikely to be identical. Recent evidence points to multiple transcriptomically-defined reactive astrocytes substates (Boghadadi et al., 2021; Garcia-Bonilla et al., 2023; Hasel et al., 2021). While these substates have not been defined at the functional level, it is most likely they play a key role in the modulation of both the inflammatory response, neuronal maintenance, and tissue preservation in the acute period after stroke.

Examining the level of astrocyte cytoskeletal elements 7- and 30 days post-stroke, TKO mice had reduced GFAP levels in astrocytes compared to WT mice. During the chronic phase of ischemic stroke, there are changes in astrocyte ramification and branch/process length (Mestriner et al., 2015; Wagner et al., 2012). A confound for the interpretation of these results is the use of a single marker, GFAP, which has increased levels following stroke – which may account for increased length and process measurements that are GFAP+ but not necessarily a change in the total number and length of processes in these astrocytes (if measured using a marker that has unaltered levels following ischemia). We have a similar caveat here.

By removing the TIC cytokines, we were able to reduce neuroinflammation during the sub-acute and chronic phases of stroke, and this translated to improved motor function in TKO mice after stroke. Functional improvement was only detected during the acute and sub-acute phases, however in most tasks mice recovered to baseline. Thus, nRAs could only play a detrimental role in early functional outcomes. However, it is more likely that this is due to the motor tasks we used and the spontaneous recovery of function in both. This is typical for these models in the chronic phase (Balkaya et al., 2013; Lai et al., 2015; Moon et al., 2009).

Recent research has found that reducing nRAs after ischemic stroke with the drug semaglutide led to changes in infarct volume, neuroinflammation, and reduction of behavioral deficits up to 28 days after stroke (Zhang et al., 2022). With the improved functional recovery of TKO mice during the acute and sub-acute phases combined with the evidence that the neuroinflammatory response and infarct volume was decreased in TKO mice during all stages of stroke, it is probable that nRAs play a detrimental role in chronic functional recovery.

Using knockout mice to examine the role of removing TIC cytokines from interacting with the neuroinflammatory response after stroke does pose some limitations. The removal of these cytokines is not just from microglia, but from all cell types throughout the whole animal, and at all developmental stages (as a non-inducible, global knockout model was used). TNF plays an important role in modulating the host defense against infections, but becomes harmful with excessive and prolonged production (Bradley, 2008). Similarly, IL-1 family members are implicated in synaptic plasticity in multiple models (Erion, 2014; Nemeth & Quan, 2021). Developmentally, C1q is integral for correct synapse pruning by microglia (Stevens et al., 2007) – a mechanism that is also integral in neurodegenerative diseases like Alzheimer’s (Hong et al., 2016). Therefore, removing these cytokines with integral physiological functions globally could introduce confounding variables as different pathways of the neuroinflammatory response are possibly being affected by the removal of these cytokines, not only the formation of nRAs.

Future research should examine potential therapeutics that target the release of these three immune factors, or intrinsic activation of nRAs, especially with clinically relevant, or potentially already approved compounds. Additional approaches include the use of astrocyte-specific conditional knockout of toxic fatty acid production enzymes, like ELOVL1, which we recently reported was protective following acute axonal injury in the optic nerve (Guttenplan et al., 2021). Further dissection of this pathway to identify how saturated long chain FFAs released from nRAs are affecting further astrocyte and microglial activation could lead to a more clinically relevant way to alter this pathway and reduce neuroinflammation after stroke. More appropriate genetic tools are already being used to investigate these pathways in other disease/trauma systems with cell-type specific and temporal control of nRA function (Guttenplan et al., 2021). Understanding this complex neuroinflammatory response following stroke would undoubtedly lead to therapies with a longer treatment window than current options.

## Acknowledgments

This work was funded by the National Institute of Health NINDS F32NS089162 to TCP, the American Heart Association Career Development Award (20CDA35310828) to TCP. NIH NEI 5R01EY033353, The Cure Alzheimer’s Fund, Alzheimer’s Association, The National Multiple Sclerosis Society, Anonymous Donors, and the Parekh Center for Interdisciplinary Neurology to SAL. Also, a Frontiers in Brain Health Award from AHA/Allen Frontiers Groups 19PABHI34580007 (MSB) and the Leducq Stroke-IMPaCT Transatlantic Network of Excellence (MSB).

